# FoxN1-dependent thymic epithelial cells promote T-cell leukemia development

**DOI:** 10.1101/247015

**Authors:** Marinella N. Ghezzo, Mónica T. Fernandes, Rui S. Machado, Ivette Pacheco-Leyva, Marta A.S. Araújo, Ravi K. Kalathur, Matthias E. Futschik, Nuno L. Alves, Nuno R. dos Santos

**Author notes:** To whom correspondence should be addressed. Nuno R. dos Santos, Instituto de Investigação e Inovação em Saúde (i3S), Rua Alfredo Allen, 208, 4200-135 Porto, Portugal, (ext.6180).

## Abstract

T-cell acute lymphoblastic leukemia (T-ALL) is an aggressive malignancy of thymocytes. The role of thymic microenvironmental cells and stromal factors in thymocyte malignant transformation and T-ALL development remains little explored. Here, using the TEL-JAK2 transgenic (TJ2-Tg) mouse model of T-ALL, which is driven by constitutive JAK/STAT signaling and characterized by the acquisition of *Notch1* mutations, we sought to identify stromal cell alterations associated with thymic leukemogenesis. Immunofluorescence analyses showed that thymic lymphomas presented epithelial areas characterized by keratin 5 and keratin 8 expression, adjacently to keratin-negative, epithelial-free areas. Both keratin-positive and -negative areas stained conspicuously with ER-TR7 (a fibroblast marker), laminin, and CD31 (an endothelial cell marker). Besides keratin 5, keratin-positive areas were also labeled by the *Ulex Europaeus* agglutinin-1 medullary thymic epithelial cell (TEC) marker. To assess whether TECs are important for T-ALL development, we generated TJ2-Tg mice heterozygous for the FoxN1 transcription factor *nude* null mutation. In contrast to *nude* homozygous mice, which lack thymus and thymocytes, heterozygous mutant mice present only mild thymocyte maturation defects. In TJ2-Tg;*Foxn1*^+/nu^ compound mice both emergence of malignant cells in pre-leukemic thymi and overt T-ALL onset were significantly delayed. Moreover, in transplantation assays leukemic cell expansion in the thymus of recipient *Foxn1*^+/nu^ mice was reduced as compared to control littermates. These results indicate that FoxN1 insufficiency impairs specifically thymic leukemogenesis but not thymocyte development.

**Summary:** In a mouse model of T-ALL, several cellular alterations were detected in thymic lymphomas, including altered epithelial distribution and increased proportion of fibroblasts or endothelial cells. Reduced dosage of FoxN1, a thymic epithelial transcription factor, delayed leukemogenesis in these mice.

## Introduction

T-cell acute lymphoblastic leukemia (T-ALL) is an aggressive hematological malignancy thought to arise from the transformation of thymocytes, as supported by similarities in the immunophenotypic, genotypic and transcriptomic profile of T-ALL cases and specific stages of intrathymic T-cell differentiation (1–3). During intrathymic thymocyte differentiation, transformation events can lead to aberrant expression of oncogenic transcription factors that induce maturation arrest (4). Genetic abnormalities in T-ALL, which include chromosomal translocations, point mutations, and deletions (4), typically result in the activation of signaling pathways commonly involved in cancer, most notably the PTEN/PI3K/AKT, Ras/MAPK and JAK/STAT pathways (5). Furthermore, one of the most common genetic alterations and considered a hallmark of T-ALL are *NOTCH1*-activating point mutations, present in more than 60% of cases (4,5). The same signaling pathways that are activated by intrinsic oncogenic mutations can also be activated by extrinsic signals provided by stromal cells in organs where T-ALL cells thrive. For example, despite the frequent presence of *Notch1* mutations, T-ALL cells were shown to respond to and be maintained *in vitro* by NOTCH ligands, such as Delta-like ligand 1 and 4 (6–8).

Several extrinsic factors promoting T-ALL growth *in vitro* and *in vivo* were identified, including interleukin (IL)-7 (9–11), IL-18 (12), insulin-like growth factor (IGF)-1 (13), ICAM-1(14) and CXCL12 (15,16). However, it remained unclear to what extent T-ALL development depends on signals from its native thymic microenvironment. In this regard, thymectomy experiments in mouse models have since long hinted that the thymic microenvironment was essential for T-cell acute leukemia/lymphoma (17,18). Early reports indicated that stromal signals are important for leukemia maintenance, as the physical contact with thymic microenvironmental cells was shown to improve survival of co-cultured thymic-derived rodent primary leukemic T cells (19,20). Furthermore, histological and immunophenotypical studies showed that the thymic stroma undergoes profound alterations throughout thymic leukemogenesis in mice (21–23). Supportive of the role of the thymic microenviroment in human T-cell leukemogenesis, several T-ALL patients present thymic enlargement at diagnosis (24,25). Moreover, patient cells were shown to expand in alymphoid fetal thymic organ cultures from NOD/SCID mouse without supplementation of exogenous factors (26).

Thymocyte development is achieved through a well-orchestrated bidirectional communication (so-called thymic crosstalk) between stromal cells and maturing thymocytes. These interactions trigger molecular changes in the thymic stromal microenvironment that are essential for thymocyte migration, differentiation and selection of functional mature T cells tolerant to self-antigens (27). The thymic microenvironment is composed of epithelial, mesenchymal, endothelial and hematopoietic cells (B lymphocytes, dendritic cells and macrophages). Thymic epithelial cells (TEC) are essential for the architecture of the embryonic and post-natal thymus, as demonstrated by genetic inactivation of the transcription factor FoxN1, an essential master regulator of TEC differentiation (28,29). The spatial organization of the thymus into compartments is associated with functionally different TEC subpopulations that are involved in the generation of successive stages of thymocyte development (30). Of note, cortical TEC (cTEC) are involved in the positive selection of thymocytes expressing functional T-cell receptor complexes, while medullary TEC (mTEC) are important for the generation of self-tolerant mature T cells. This function is mediated through MHC class II-mediated presentation of tissue-restricted self-antigens, which are essential for the negative selection of auto-reactive thymocytes and the generation of regulatory T cells (31).

The thymic molecular and cellular players involved in stromal support of thymocyte leukemogenesis have only recently began to be explored. Our group has reported that stromal cell inactivation of the RelB transcription factor and the TNF receptor superfamily member lymphotoxin-β receptor (LTβR), both components of the noncanonical NF-κB signaling pathway, delayed leukemia development in the TEL-JAK2 transgenic (TJ2-Tg) mouse model (32,33). Recently, Triplett and colleagues (34) found in another T-ALL mouse model that IGF1R was highly expressed by malignant cells and that high IGF-1 secretion by tumor-associated thymic dendritic cells (DCs) favored leukemic cell survival. Together, these data suggest that thymic microenvironmental cells participate in the development of T-cell leukemia.

In the present study, we characterized the thymic stromal alterations at various stages of disease and found that thymic leukemogenesis was associated with an increase in mTEC but not cTEC molecular markers. More importantly, we found that haploinsufficiency of the *Foxn1* gene delayed thymic leukemogenesis.

## Materials and methods

### Mice

Mice were bred and maintained at the CBMR/University of Algarve barrier animal facility (HEPA filtration of incoming air, differential pressure, and disinfection or sterilization of room equipment and supplies), under 12 h light/dark cycles, and with autoclaved food (4RF25 diet; Mucedola, Settimo Milanese, Italy) and water *ad libitum*. Microorganism screening (Charles River, L’Arbresle, France and Idexx Bioresearch, Ludwigsburg, Germany) detected opportunistic pathogens (*Helicobacter* spp., *Pasteurella pneumotropica* and murine norovirus) in the housing room. The animal house and project were approved by Portuguese authorities (*Direção-Geral de Agricultura e Veterinária*). All experimental procedures were performed in accordance with European (Directive 2010/63/UE) and Portuguese (*Decreto-Lei* nº113/2013) legislation. EµSRα-TEL-JAK2 (Tg[Emu-ETV6/JAK2]71Ghy) transgenic mice were described before (33,35). *Foxn1*^*+/nu*^ (B6.Cg-*Foxn1*^*nu*^/J) mice, which carry the *nude* (*nu*) null allele, were obtained from *Instituto Gulbenkian de Ciência* (Oeiras, Portugal). Upon breeding the two strains, always on the C57BL/6 genetic background, we obtained TJ2-Tg;*Foxn1*^+/nu^ but no viable *nu*/*nu* mice. *Foxn1*^*+/nu*^ mice were PCR genotyped using primers 5’-GGC CCA GCA GGC AGC CCA AG and 5’-AGG GAT CTC CTC AAA GGC TTC CAG, followed by *Bsa*JI PCR product digestion. For transplantation experiments, thymic-derived leukemic cells were prepared from diseased TJ2-Tg mice, suspended in phosphate-buffered saline (PBS) solution and injected into the tail vein of non-anesthetized 8 to 12 week-old wild-type (WT) C57BL/6 mice or into 6 to 9 week-old *Foxn1*^+/+^ and *Foxn1*^+/nu^ littermate mice (4 × 10^6^ cells per mouse). Mouse sex is indicated in figures. TJ2-Tg and leukemic cell recipient mice were monitored for leukemia development, and euthanized by CO_2_ inhalation when manifesting signs of disease and reaching pre-defined endpoints (e.g. dyspnea, perceptible loss of activity, enlargement of lymph nodes and/or abdomen).

### Flow Cytometry

Single-cell suspensions from collected organs were prepared in PBS by tissue dissociation against a 70 µm cell strainer (BD Biosciences, San Jose, CA, USA). Cells were collected by centrifugation and resuspended in staining buffer (PBS with 10% fetal bovine serum [FBS; PAA Laboratories, Linz, Austria] and 10 mM NaN_3_) with fluorochrome-labeled antibodies. After washing with staining buffer, cells were analyzed in a FACS Calibur flow cytometer (BD Biosciences). Fluorescein isothiocyanate (FITC)-, R-phycoerythrin (PE)-, PE-cyanine 5 (PE-Cy5)-, or Allophycocyanin (APC)-conjugated antibodies specific for CD25 (PC61), CD4 (GK1.5), CD8 (53-6.7), and CD44 (IM7; BioLegend, San Diego, CA, USA) were used. Propidium iodide (P4170, Sigma-Aldrich, St. Louis, MO, USA) staining was used to gate viable cells for analysis. Analyses were performed on CellQuest (BD Biosciences) and FlowJo (FlowJo LLC, Ashland, OR, USA) software.

### Immunofluorescence

Whole thymi were included and frozen in OCT compound (VWR, Radnor, PA, USA). Thymic cryosections of 6-10 µm were dried at 37ºC for 30 min before fixation with pre-cold acetone for 10 min at room temperature. Sections were re-hydrated in PBS, washed with PBSW solution (PBS with 0.1% Tween 20 [VWR]) and incubated with blocking solution (10% BSA [A9418, Sigma-Aldrich], 20% FBS in PBSW) for 1 h. Primary antibodies (Supplementary Table 1) were incubated overnight at 4ºC, washed with PBSW, followed by 1 h incubation with secondary antibodies (Supplementary Table 1) at room temperature. Slides were washed with PBS and mounted in Mowiol reagent with 0.15% DAPI (Biotium, Fremont, CA, USA). Images were acquired with an Axio Imager Z2 fluorescence microscope (Carl Zeiss, Jena, Germany). Images were processed using open-source ImageJ and AxioVision (Carl Zeiss) software. Assembly of sequential images was performed using Pairwise Stitching in Fiji software.

### Quantitative and semi-quantitative RT-PCR

Stromal cell-enriched fractions and thymocyte suspensions were obtained by mechanically dissociating thymic tissue through a 70 µm cell strainer in PBS. Total RNA from whole thymus and stromal cell- or thymocyte-enriched thymic samples was prepared using Trizol reagent (Life Technologies, Carlsbad, CA, USA) and phenol:chloroform extraction, according to the manufacturer’s instructions. Genomic DNA was removed from RNA samples with DNase I treatment (Thermo Fisher Scientific, Waltham, MA, USA). Reverse transcription was performed using the First Strand cDNA Synthesis Kit (Thermo Fisher Scientific) and oligo(dT)_18_ primers, according to the manufacturer’s instructions. Semi-quantitative PCR was performed in 25 µL volume including 1X GoTaq Flexi Buffer, 1.5 mM MgCl_2_, 0.2 mM dNTPs, 0.1 mM of each primer (Supplementary Table 2), and 0.5 U of GoTaq DNA Polymerase (Promega, Madison, WI, USA). Serial dilutions (1/20, 1/60, and 1/180) of template cDNA was used for each sample. PCR reactions were performed in a C1000 thermal cycler (BioRad, Hercules, CA, USA) according to the following protocol: initial denaturation at 94ºC for 5 min, 26 to 35 cycles of 94ºC for 1 min, 60ºC for 1 min, and 72ºC for 1 min, and a final extension at 72ºC for 7 min. For quantitative PCR, 2 µL of 1/20 diluted cDNA, SsoFast EvaGreen Supermix (Bio-Rad) and 300 nM of primers (Supplementary Table 2) in 20 µL were analyzed on a CFX 96 Real-time PCR detection system (Bio-Rad). The mean fold change in expression of the gene of interest was calculated in relation to expression of a reference gene (*Hprt*) by the comparative C_T_ method (2^-∆∆C(T)^ method).

### Genomic DNA extraction and Notch1 mutation detection

Genomic DNA was extracted from TJ2-Tg leukemic cells using the GeneJET Genomic DNA purification kit (Thermo Fisher Scientific), following the manufacturer’s protocol. The exon 34 of *Notch1* was PCR amplified as previously described (36), and purified PCR products were sequenced at the CCMAR-Centre of Marine Sciences sequencing facility (Faro, Portugal).

### Next Generation Sequencing

Thymic stromal cell-enriched fractions from WT (5 male) and TJ2-Tg (1 female and 2 male) mice were obtained as described above and total RNA was extracted using the Direct-zol RNA Miniprep kit (Zymo Research). Next, mRNA was isolated using the NEBNext Poly(A) mRNA Magnetic Isolation Module (New England Biolabs, Ipswich, MA, USA), following the manufacturer’s instructions. The WT and TJ2-Tg mRNA samples were pooled and purified using the RNeasy MinElute Cleanup kit (Qiagen) and further processed using the Ion Total RNA-Seq kit v2 (Thermo Fisher Scientific). Sequencing was performed on Ion Torrent PGM (Thermo Fisher Scientific) using Ion 316 Chip kits v2 (Thermo Fisher Scientific) for the WT and TJ2-Tg pooled samples. Both FastQC and Prinseq indicated high sequence quality except for a residual adapter sequence in a subpopulation of reads. The latter was removed by Cutadapt software. Additionally, reads were trimmed when the corresponding Phred score fell below 25. Subsequently, reads were aligned to the full mouse genome (UCSC, version mm10) by the Torrent Mapping Alignment Program (TMAP). In total, 3,204,955 WT and 3,125,039 TJ2-Tg reads were mapped to the mouse genome. Gene expression levels were obtained by HTseq-count, and differential gene expression was assessed with Bioconductor package edgeR.

### Statistical analysis

Graphs and statistical analyses were performed with GraphPad Prism software (La Jolla CA, USA). Statistical tests are indicated in the figure legends. Two-tailed *t*-tests were used to determine how significant were the differences between the means of two independent samples. Welch’s correction was applied when variances were unequal, as determined by the F-test.

## Results

### TEL-JAK2 mouse thymic lymphomas present profound structural changes as compared to healthy thymi

Thymic lymphomas from TJ2-Tg mice often become 10-20 times larger than age-matched control thymi and lose the typical thymic compartmentalization in cortex and medulla (35,37). To determine whether the thymic microenvironment influences TJ2-Tg-induced leukemogenesis, we began by assessing the spatial distribution of several thymic stromal cells in diseased mice.

Immunofluorescence analysis of thymic cryosections with antibodies against the keratin (Krt) 5 and Krt8 epithelial cell markers revealed large cellular regions that lack Krt5- and Krt8-labeling, here designated as keratin-negative areas (KNA) (Supplementary Figure 1A and B). In contrast to the WT thymus, where medullary (Krt5-positive) and cortical (Krt8-positive, Krt5-negative) regions were clearly spatially segregated (Figure 1, panels a1 and a2), keratin-positive areas in most TJ2-Tg thymic lymphomas displayed intermingled or overlapping Krt5 and Krt8 expression (Figure 1, panels i1 and i2, and Supplementary Figure 1C). Further spatial characterization based on Ulex Europaeus agglutinin-1 (UEA-1) labeling and DEC205 expression, which respectively identify mTECs and cTECs (Figure 1, panels b1-b2, and c1-c2), showed that UEA-1^+^ mTECs were found within Krt5^+^ areas of TJ2-Tg thymic lymphomas, (Figure 1, panels j1 and j2).

**Fig.1.**
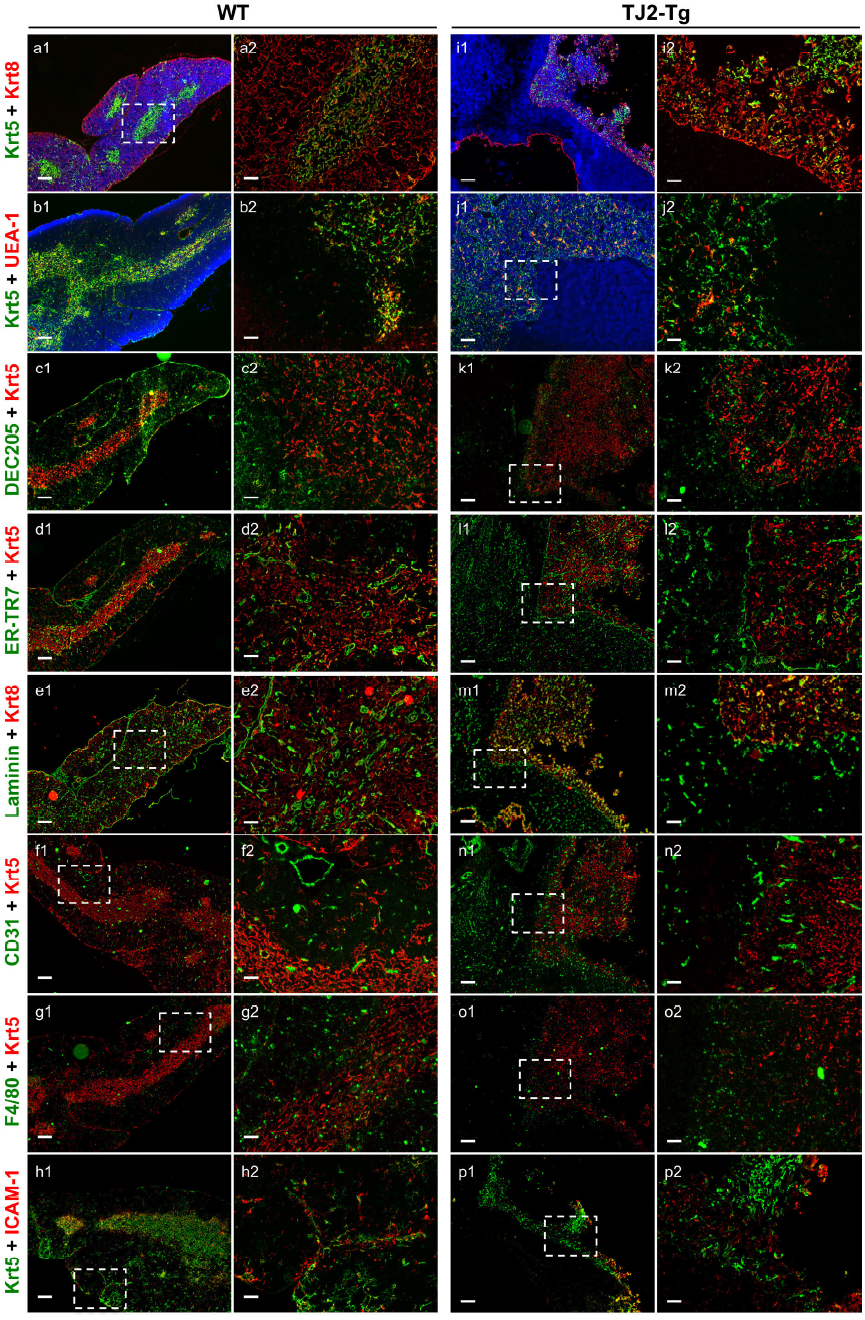
Characterization of the thymic stromal cellular alterations in advanced stage thymic lymphomas from TJ2-Tg mice. Double immunofluorescence staining of a WT thymus (panels a1-h2) and two representative TJ2-Tg thymic lymphoma (mouse nº 341, panels i1-i2, k1-p2; mouse nº 128, panels j1-j2; out of at least three analyzed) with antibodies against the indicated markers. The higher magnification panels depict a thymic region encompassing both medulla and cortex (panels a2-g2) and keratin-positive and negative boundaries (panels i2-p2). For panels a1, b1, i1 and j1 DAPI staining (blue) is shown. Scale bars: 200 µm (panels a1-p1), 400 µm (panels a2-p2). When applicable, dashed-line rectangles represent the magnified area.

DEC205 (encoded by the Ly75 gene) is also a marker of thymic DCs (38). TJ2-Tg2 thymic lymphomas displayed a dense DEC205 staining in keratin-positive regions, notably stronger in areas with weaker Krt5 staining, suggesting that DEC205 labels residual cortical areas marked by ectopic Krt5 expression (Figure 1, panels k1 and k2). Interestingly, DEC205^+^ cells were found within thymic lymphoma KNAs (Figure 1, panel k1 and k2), indicating the presence of DCs.

As reported, fibroblast labeling by the ER-TR7 antibody was not restricted to any particular region of the WT thymus, but was more abundant in the thymic capsule, trabeculae, and lining of blood vessels (Figure 1, panels d1 and d2) (39). In TJ2-Tg thymic lymphomas, the pattern of ER-TR7 expression was similar in both keratin-positive and -negative areas, although a layer of ER-TR7^+^ cells marked the border between these two areas (Figure 1, panels l1 and l2, and Supplementary Figure 2A). The expression pattern of laminin labeling was similar to the ER-TR7 labeling, in that it was present throughout the tumor tissue, irrespective of keratin labeling, and bordering keratin-positive and - negative areas (Figure 1, panels m1 and m2). The CD31 marker, labeling endothelial cells, was also present throughout the thymic lymphoma tissue (Figure 1, panels n1 and n2). Double immunofluorescence analyses showed that all CD31^+^ lymphoma blood vessels were surrounded by laminin (Supplementary Figure 2B), which is normally a component of vascular basement membranes (40).

While F4/80^+^ macrophages were scattered in normal thymic tissue (Figure 1, panels g1 and g2), these cells were also distributed throughout TJ2-Tg thymic lymphomas, albeit more abundantly in keratin-positive areas (Figure 1, panels o1 and o2). Finally, we also analyzed the distribution pattern of the integrin-binding ICAM-1 adhesion molecule. As previously reported (41,42), ICAM-1 was detected both in the thymus cortex and medulla, although more prominently in the latter (Figure 1, panel h1). In TJ2-Tg thymic lymphomas, ICAM-1 expression was more prevalent in areas of stronger Krt5 labeling (Figure 1, panels p1 and p2), often co-localizing at the cellular level (Supplementary Figure 2C). These data indicate that thymic lymphoma mTEC-like cells, like normal mTECs, express ICAM-1.

The immunohistological analyses indicated that TEC subpopulations undergo profound alterations in spatial distribution and organization upon lymphoma development. Gene expression analysis of whole TJ2-Tg thymic lymphomas revealed a significant decrease of Krt5 and Krt8 transcripts as compared to normal whole thymi, providing further indirect evidence that the proportion of epithelial cells within lymphomas was reduced (Supplementary Figure 3). Furthermore, the ratio of Krt5/Krt8 mRNA levels was significantly higher in TJ2-Tg thymic lymphomas than in normal thymi (Supplementary Figure 3), thus supporting the notion that the mTEC/cTEC ratio is enhanced in those tumors.

To gain insights into the dynamics of stromal cell alterations during TJ2-Tg disease, we performed immunohistology on smaller TJ2-Tg thymic lymphomas, i.e. weighing ≤240 mg (mean of 166 mg; n=4) as opposed to ≥589 mg (mean of 903 mg; n=4) described above. These analyses led to two main findings: first, Krt5 labeling was expanded in relation to Krt8 labeling, with concomitant loss of the normal cortico-medullary compartmentalization (Figure 2A, second row panels); second, the extension of KNAs, which contained abundant ER-TR7^+^ fibroblasts, increased with lymphoma size (Figure 2A, third row panels). Since an evident expansion of Krt5^+^ in relation to Krt8^+^ areas was recurrently detected in TJ2-Tg thymic lymphomas, we have also immunostained thymic lymphomas for the UEA-1 medullary marker, and this led to a similar staining pattern as Krt5 (Figure 2A, bottom panels). These results indicate that in small lymphomas Krt5^+^UEA-1^+^ TECs expanded in detriment of Krt8^+^ TECs.

**Fig.2.**
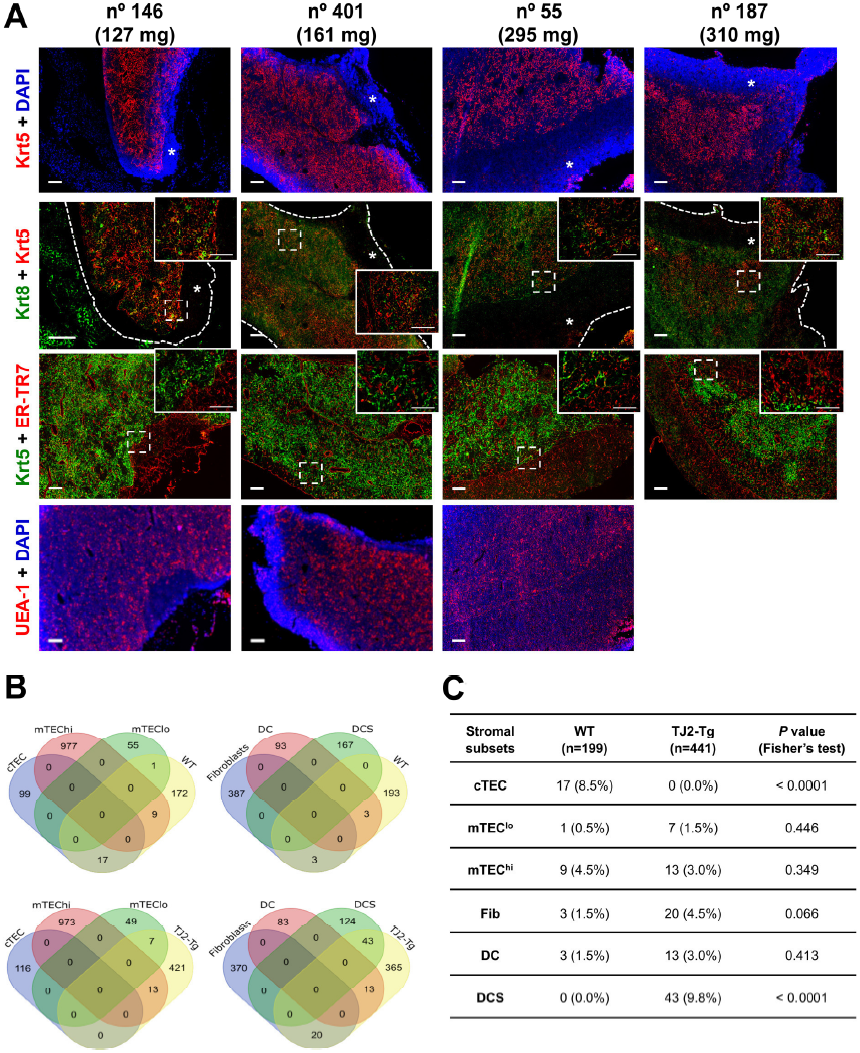
Early stages of TJ2-Tg lymphomagenesis are accompanied by thymic medullary expansion and cortical reduction. Cryosections from TJ2-Tg thymic lymphomas of the indicated weights were immunostained for Krt5, Krt8, and UEA-1 thymic epithelial markers and the ER-TR7 fibroblast marker. Asterisks indicate KNAs; dashed-line rectangles represent area magnified in the inset panel. Scale bars: 200 µm (main panels), 100 µm (inset panels). **(B)** Venn diagrams displaying the overlap between gene sets characteristic for cortical thymic epithelial cells (cTEC), medullary thymic epithelial cells (mTEChi and mTEClo), dendritic cells (DC and DCS, negative and positive for Sirpα, respectively) and fibroblasts from the Ki et al. study (43) and upregulated genes in WT versus TJ2-Tg (T-ALL) thymic stromal fractions. (C) For each stromal cell subset, the number and percentage of genes differentially expressed between WT versus TJ2-Tg (T-ALL) thymic stromal fractions is given. *P* values determined by two-tailed Fisher’s exact test.

### Global gene expression analysis reveals alterations in thymic stromal cell composition early in leukemogenesis

To gain further insights into the nature of the earliest stromal cell alterations in thymic leukemogenesis, we used a genome-wide transcriptomic approach. To this end, stromal cell-enriched thymic fractions were obtained from 8- to 10-week-old WT mice and TJ2-Tg mice with incipient disease, as defined by nearly normal sized thymi (<100 mg) and presence of low proportions (10-22%) of CD8^+^CD25^+^ leukemic cells within the thymocyte population (not shown). Pilot experiments showed that the crude fractionation method applied led to enrichment of thymic stromal cell genes and depletion of thymocyte-specific genes in stroma-enriched fractions (Supplementary Figure 4). Upon RNA extraction, pooled samples were subjected to high-throughput sequencing. Bioinformatic analyses disclosed 640 differentially expressed genes (DEGs), 199 upregulated in WT thymi (Supplementary Table 3) and 441 upregulated in TJ2-Tg thymi (Supplementary Table 4). As expected from the crude stromal fractionation, several genes differentially expressed between WT and TJ2-Tg samples most likely derive from non-excluded T lineage cells (e.g., *Ptcra*, *Rag1*, *Rag2*, *Dntt*, *Themis*, etc.). Nevertheless, many up- or downregulated genes were well-characterized stromal cell markers (e.g. *Ccl25*, *Vim*, *Lepr*, *S100a4*, *Zbtb46*, *Xcl1*, etc.). To know if the identified DEGs reflect imbalances in thymic stromal cell proportions, we investigated whether they matched genes specifically and uniquely expressed in particular thymic stromal cell subsets, as identified by Ki and colleagues (43). Interestingly, of 116 cTEC-specific genes, 17 were upregulated in WT compared to TJ2-Tg thymi whereas none was upregulated in TJ2-Tg compared to WT thymi (Figure 2B and C, and Supplementary Table 5). These findings confirm that cTEC markers are downregulated during lymphomagenesis. In contrast, mTEC-specific genes were not significantly enriched in either WT or TJ2-Tg samples. In addition, we observed that several genes specific of Sirpα^+^ DCs were upregulated in TJ2-Tg but not WT thymi (Figure 2B and C and Supplementary Table 5). These data indicate that thymic leukemogenesis in TJ2-Tg mice is accompanied by alterations in TEC subset proportions and an increase in DCs.

### Foxn1 *haploinsufficiency delays the onset of TEL-JAK2-induced leukemia*

To assess whether TECs play a role in thymic leukemogenesis, we generated TJ2-Tg mice lacking one *Foxn1* allele, a gene crucial for TEC development and maintenance in a dose-dependent manner (28). Mice carrying the *Foxn1 nude* mutation in heterozygosity presented normal proportions of thymocyte developmental stages (as defined by CD4, CD8, CD25 and CD44 expression), including those ranging from the CD4^−^CD8^−^CD25^+^CD44^−^ double negative three (DN3) stage to the CD4^+^CD8^+^ double-positive stage (Supplementary Figure 5A and B), which are the main cellular targets for TJ2-Tg malignant transformation (37). It was reported that pre-puberty *Foxn1*^+/nu^ mice present lower thymic weight than WT controls (28,44,45). Since TJ2-Tg leukemia arises during adulthood, we analyzed older mice. Thus, we observed that 8 or 20 week-old *Foxn1*^+/nu^ mice presented slightly smaller thymi than *Foxn1*^+/+^ littermates (1.1-1.3-fold difference) (Supplementary Figure 5B), but no statistically significant differences in thymic weight were found when considering females alone or older mice (7- to 9-months) (Supplementary Figure 5C and D). Strikingly, comparison of *Foxn1*^+/+^ and *Foxn1*^+/nu^ TJ2-Tg mouse cohorts showed that the onset of leukemia was delayed in TJ2-Tg mice with *Foxn1* haploinsufficiency (7-week difference in median survival; hazard ratio of 2.93, with 95% CI of 1.90 to 4.51; Figure 3A). Both TJ2-Tg;*Foxn1* cohorts included both sexes, with an equal male-to-female ratio of 1.6:1. Survival analysis of each sex separately revealed that *Foxn1* haploinsufficiency delayed leukemia onset in both settings (Figure 3B). Despite the different survival rates of the two cohorts, no significant differences were observed regarding end-stage tumor load (Figure 3C). Moreover, end-stage TJ2-Tg;*Foxn1*^+/nu^ thymic lymphomas displayed KNAs and areas of overlapping Krt5 and Krt8 staining as control lymphomas (Supplementary Figure 6). Like in other T-ALL mouse models, TJ2-Tg leukemic cells were found to acquire activating frameshift mutations in the Notch1 exon 34, encoding the PEST domain (>65% incidence) (Supplementary Table 6). Notch1 PEST domain heterozygous mutations, as illustrated by the simultaneous detection of WT and mutated sequences (Supplementary Figure 7), were also detected, with similar frequency, in leukemic cells from both *Foxn1* haploinsufficient and control TJ2-Tg mice (Table 1). These mutations consisted of deletions and/or insertions, often affecting the previously reported exon 34 hotspots (codons 2361 and 2398/99) (46,47) and leading to premature stop codons and predicted loss of the C-terminal PEST domain (Table 1). Together, these data indicate that FoxN1 insufficiency delayed leukemogenesis without influencing the outcome of disease evolution.

**Table 1.**
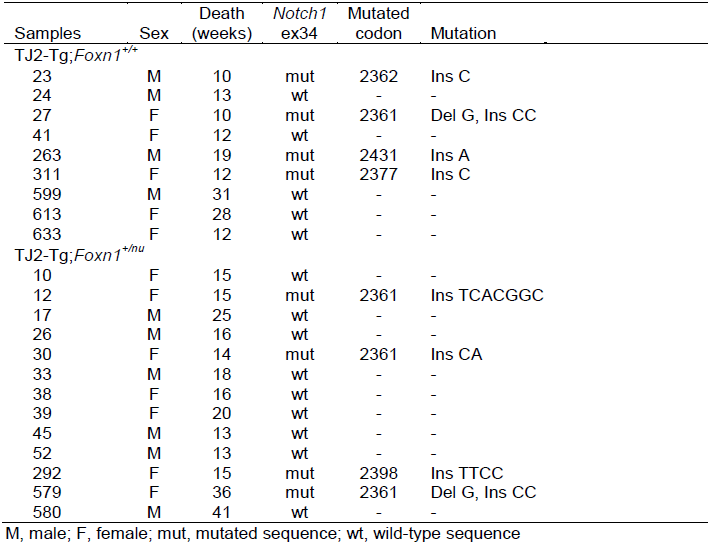
Exon 34 *Notch1* mutations in TJ2-Tg-induced mouse leukemia samples with distinct *Foxn1* genetic backgrounds.

**Fig.3.**
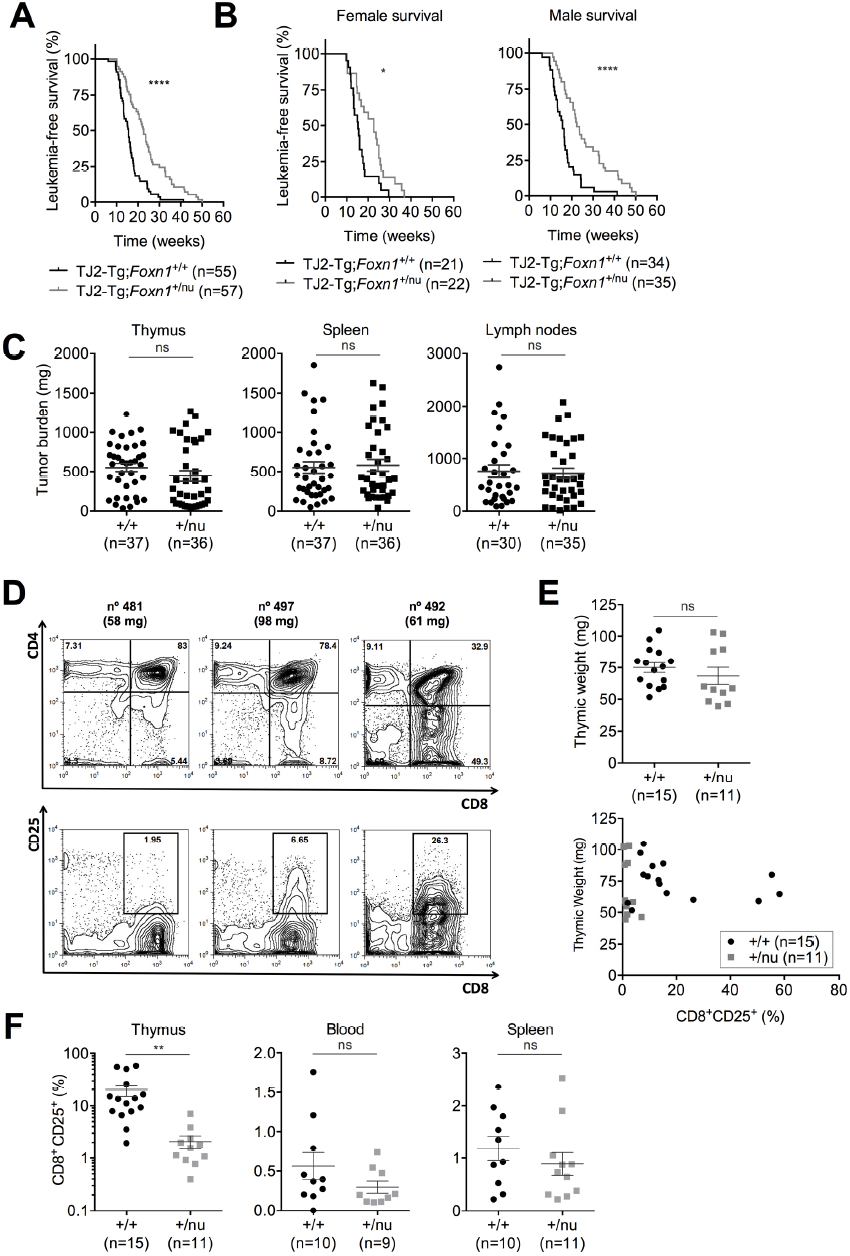
Foxn1 haploinsufficiency delays TEL-JAK2-induced leukemogenesis. **(A)** Kaplan-Meier leukemia-free survival curves for cohorts of TJ2-Tg;*Foxn1*^+/+^ and TJ2-Tg;*Foxn1*^*+/nu*^ mice, with median survivals of 16 and 23 weeks, respectively (log-rank test, *P* value ≤ 0.0001). Both groups of mice include an equal ratio (1:1.6) of females to males. **(B)** Kaplan-Meier leukemia-free survival curves for cohorts of female or male mice. The median survival of females was 15 and 23 weeks for TJ2- Tg;*Foxn1*^+/+^ and TJ2-Tg;*Foxn1*^+/nu^, respectively (log-rank test, *P* value ≤ 0.05), and of males was 16 and 22 weeks for TJ2- Tg;*Foxn1*^+/+^ and TJ2-Tg;*Foxn1*^+/nu^, respectively (log-rank test, *P* value ≤ 0.0001). **(C)** Weights of thymus, spleen and lymph nodes collected from full-blown diseased TJ2-Tg;*Foxn1*^+/+^ and TJ2-Tg;*Foxn1*^*+/nu*^ mice. **(D)** Thymocyte flow cytometry analysis of three representative 8-week-old healthy TJ2-Tg mice (thymic weight of each mouse is indicated) with different proportions of aberrant CD8 single-positive (top panels) and CD8^+^CD25^+^ cells (bottom panels). Percentage of cells in each region is shown. **(E)** Thymic weight of 8 week-old TJ2-Tg;*Foxn1*^+/+^ and TJ2-Tg;*Foxn1*^*+/nu*^ littermate mice (top panel) and its correlation with percentage of detected CD8^+^CD25^+^ thymocytes (bottom panel). **(F)** Percentage of CD8^+^CD25^+^ leukemic cells in thymic, splenic, and blood cell suspensions from same mice as in (E). (C, E, F) Data are presented with mean ± SEM. *P* values determined by two-tailed, unpaired *t*-test with Welch’s correction (**, *P* ≤ 0.01; *ns*, non-significant).

To investigate whether *Foxn1* haploinsufficiency delayed the early stages of leukemogenesis at the cellular level, we monitored disease emergence in young pre-disease TJ2-Tg mice through detection of CD8^+^CD25^+^ leukemic cells within total thymocytes, (Figure 3D). At this age, leukemic cells can be detected in TJ2-Tg thymi without significant organ enlargement (33). Indeed, thymic weight in the analyzed *Foxn1*^+/+^ and *Foxn1*^+/nu^ TJ2-Tg mice was not significantly different and it did not correlate with the percentage of detected CD8^+^CD25^+^ leukemic cells (Figure 3E). However, thymi from TJ2-Tg;*Foxn1*^+/nu^ mice displayed a significantly lower percentage of CD8^+^CD25^+^ cells than TJ2-Tg;*Foxn1*^+/+^ thymi (7.8-fold difference; Figure 3F). At this age, leukemic cells are detectable mainly in the thymus (33). Although low percentages of CD8^+^CD25^+^ cells could be detected in the blood, bone marrow and spleen of some mice, those were not significantly different among *Foxn1* genotypes (Figure 3F and not shown). These findings indicate that leukemic cell emergence is delayed when *Foxn1* dose is reduced in TEC.

Although *Foxn1* haploinsufficiency leads to a mild decrease in thymic size, which could contribute to the observed delayed leukemogenesis, the above-presented data suggest that reduction in FoxN1 expression levels in TECs has a more marked impact on leukemogenesis than on thymocyte generation and differentiation.

### *The* Foxn1-*dependent thymic microenvironment facilitates leukemic cell expansion*

To determine the role of the *Foxn1*-dependent microenvironment in leukemia maintenance, leukemic cells were collected from diseased TJ2-Tg mice and infused into nonconditioned syngeneic *Foxn1*^+/nu^ and *Foxn1*^+/+^ littermate controls (Figure 4A). When manifesting signs of disease, recipient mice were killed for macroscopic and flow cytometry analyses. Thymi from all recipient mice became infiltrated by CD8^+^CD25^+^ leukemic cells (Figure 4B). Two independent experiments using either male or female recipients were performed and each individually revealed a trend towards lower tumor load in *Foxn1*^+/nu^ thymi (Figure 4C). Following normalization and data pooling, we found that *Foxn1*^+/nu^ mice displayed statistically significant lower thymic weight than *Foxn1*^+/+^ littermates (Figure 4D). In contrast, the relative splenic and hepatic tumor load was not influenced by *Foxn1* genotype (Figure 4D). These results thus indicate that *Foxn1* expression in TECs promotes leukemic cell expansion in the thymus.

**Fig.4.**
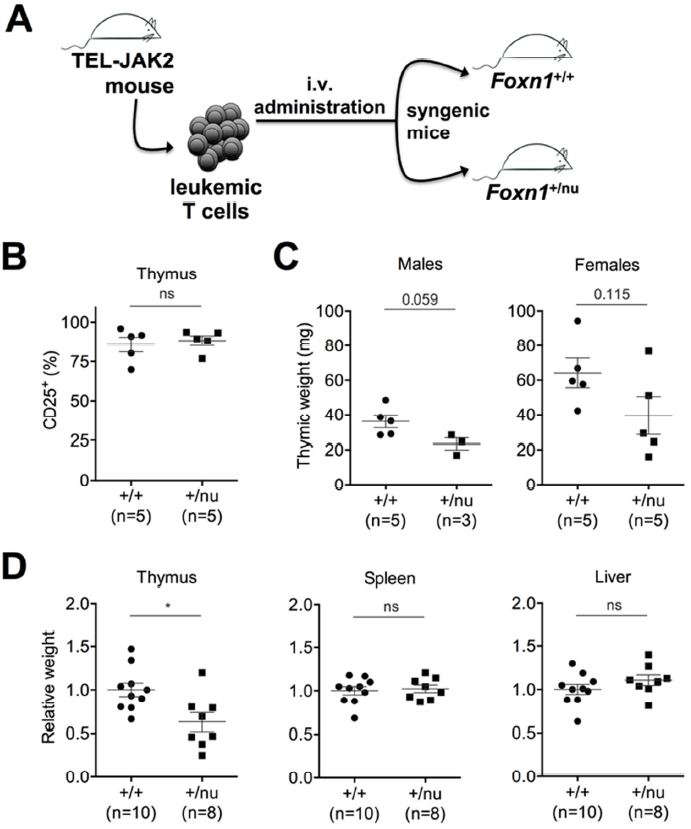
*Foxn1* haploinsufficiency impairs leukemic cell expansion in recipient mouse thymi. **(A)** Transplantation experiment design. **(B)** Flow cytometry detection of CD25^+^ leukemic cells in thymic cell suspensions of recipient littermate mice with the indicated genotype. Data shown is representative of one of two independent experiments performed with leukemic cells from two different TJ2-Tg mice. **(C)** Thymic weights of recipient mice from two independent experiments, one with male and another with female recipients. **(D)** Relative organ weights from recipient mice of transplantation experiments shown in (C) pooled after normalization based on the average weight of the WT (*Foxn1*^+/+^) animals in each experiment. (B-D) Data are presented with mean ± SEM; *P* value determined by two-tailed, unpaired *t-*test (*, *P* ≤ 0.05; *ns*, non-significant).

## Discussion

The malignant transformation of mouse or human thymocytes is triggered by the deregulation of key cellular mechanisms due to both cell-intrinsic genetic alterations and microenvironmental stimulation (48,49). Here, using a mouse model of JAK kinase-driven and *Notch1*-mutated T-ALL, we identified several microenvironmental cellular alterations associated with thymic lymphoma development and progression. These included shifts in expression of TEC markers, with loss of cortical and increase of medullary epithelial markers, together with increased proportions of fibroblasts, DCs and endothelial cells. Highlighting a role for TECs, we found that reducing the genetic dose of FoxN1, an essential TEC transcription factor in TEC differentiation, delayed thymic leukemogenesis.

Diseased TJ2-Tg mice present large thymic lymphomas with loss of cortico-medullary demarcation and frequent disruption of the thymic capsule (32,35). By performing immunohistological analyses against thymic keratin proteins, we observed that TJ2-Tg mouse thymic lymphomas developed conspicuous KNAs, which were more extensive in larger lymphomas. Similar observations were reported in thymic lymphomas from other mouse models (21,34). In TJ2-Tg mice, KNAs included fibroblasts and laminin-containing extracellular matrix, suggesting that leukemic cells first invade the capsular connective tissue before expanding outwardly, thus generating lymphomatous KNAs. By studying spontaneous thymic lymphomas of different sizes, we found that the normal cortico-medullary demarcation was always lost. This phenomenon was accompanied by a strong reduction in cortical (Krt8-positive, Krt5-negative) areas, which is in agreement with earlier reports of cortical atrophy and medullary expansion at pre-leukemic stages of viral-induced murine thymic lymphoma (22,23,50–52). Our data thus support the concept that upon emergence of leukemic cells in the thymus, these disseminate throughout all thymic compartments, impacting on TEC differentiation and/or distribution, before invading the thymic capsule and undergoing further expansion.

FoxN1 is a pivotal transcription factor for thymus and skin epithelial cell differentiation in rodents and humans (53). However, its role in cancer is less understood. Previous evidence indicated that FoxN1 suppresses the transition from benign to malignant keratinocyte lesions (54). Another report supporting a tumor-suppressive role for *Foxn1* showed that its haploinsufficiency accelerated thymic leukemogenesis in the endogenous retrovirus-expressing AKR murine strain (55). In contrast with these data, we found that loss of a single *Foxn1* allele hindered both the onset of spontaneous TEL-JAK2-induced leukemia/lymphoma and the thymic expansion of leukemic cells in recipient mice. Since leukemogenesis in TJ2-Tg mice is initiated in the thymus (33,37) and only TECs express *Foxn1*, we conclude that the delayed leukemogenesis in heterozygotes is caused by FoxN1-mediated defects dependent on these cells. *Foxn1* haploinsufficiency was shown to cause a moderate reduction in thymic weight of pre-puberty mice (under 5 weeks) of different outbred and inbred strains (28,44,45). Young adult *Foxn1*^*+/nu*^ mice (8 and 20 weeks of age) displayed a mild reduction in thymic weight, which was less evident considering females alone or older mice, but no defects in the major *Foxn1*^+/nu^ thymocyte subsets were apparent. Further supporting the notion that delayed leukemogenesis caused by *Foxn1* haploinsufficiency is not due to a mere decrease in thymocyte cellularity, the decrease in adult thymic weight caused by loss of one *Foxn1* allele was much less pronounced (1.3-fold) relatively to the delay in both emergence of thymic leukemic cells (7.8-fold) and leukemia-related TJ2-Tg mouse death (2.2-fold higher mortality rate at 20 weeks of age). Although a putative role for reduced thymocyte cellularity cannot be formally excluded, our data indicate that the delayed thymic leukemogenesis caused by *Foxn1* haploinsufficiency results from an as yet unidentified cellular mechanism acting specifically on leukemic cell propagation but not on thymocyte development.

Although the FoxN1-mediated mechanism fostering leukemogenesis remains unclear, it is possible that leukemic cell development is more sensitive than thymocyte development to changes in expression of specific FoxN1 target genes. In this line, we observed that *Foxn1* haploinsufficiency was associated with a mild reduction in Ccl25 thymic expression (not shown), a well-established FoxN1 target gene (29,56) and a potential T-ALL growth factor (57). Future studies should address whether this or other FoxN1 target genes (e.g. *Dll4* and *Cxcl12*) play direct roles in thymic leukemogenesis. Alternatively, FoxN1 could impinge on leukemogenesis indirectly through changes in biological properties of TECs or other thymic microenvironmental cells. Since thymic leukemogenesis is accompanied by a disorganization in TEC marker expression, it is possible that *Foxn1*^*+/nu*^-associated TEC differentiation defects underlie the delayed leukemia onset in TJ2-Tg;*Foxn1*^*+/nu*^ mice. In addition, TECs were shown to regulate thymic vascularization (58,59), and thymic DC development, differentiation and recruitment (60–62). Whether *Foxn1* haploinsufficiency in TECs is linked to vascular or DC defects warrants further investigation.

It was previously shown that RelB and LTβR proteins, both expressed in mTEC and other thymic stromal cells, contribute to leukemogenesis (32,33). Additionally, these proteins are essential for thymic medulla development and maintenance, as shown by the impaired medullary compartment of the respective knock-out mice (63,64). The notion that RelB, LTβR and FoxN1 hinder leukemogenesis and that all are expressed in mTECs suggest that these or other medullary cell types are involved in thymic leukemogenesis.

Using a panel of thymic cell markers, we detected major organizational alterations involving several thymic stromal cell types at early and advanced stages of TJ2-induced leukemia. Most notably, we observed increased expression of CD31 and ER-TR7, markers for endothelial cells and thymic fibroblasts, respectively, suggesting that angiogenesis, fibroblast proliferation and accrued ECM deposition, which are often associated with cancer (65) may favor thymic lymphomagenesis, especially at the most advanced stages. We have also found expression of markers for macrophages (F4/80) and DCs (DEC205) in lymphomas. Macrophages have been identified in mouse thymic lymphomas (22,51,66), and recent studies indicate that lymphoma-associated macrophages stimulate T-ALL proliferation *in vitro* (67). Thus, the potential involvement of tumor-associated macrophages in T-cell leukemogenesis merits further investigation. DCs are usually associated with an anti-tumoral role via their function in adaptive immunity as tumor antigen-presenting cells (68). However, it was recently shown that DCs favored survival or proliferation of B and T leukemic cells (34,69). More specifically, Triplett et al. (34) reported that thymic lymphoma-derived DCs (most notably Sirpα-positive DCs) promoted leukemic T cell survival through IGF-1 and PDGF (platelet-derived growth factor) secretion. An increased expression of Sirpα^+^ DC genes was found in the stroma of TJ2-Tg thymic lymphomas, so studies determining whether DCs are involved in TJ2 leukemogenesis are warranted.

Together, our findings demonstrate that the thymic microenvironment in general and TECs specifically are crucial for the malignant transformation of thymocytes. The identification of the key molecular signals produced by stromal cells will not only shed further light on the mechanisms underlying T-cell leukemogenesis, but may also be instrumental for the design of therapies targeting these malignancies.

## Acknowledgements

We thank James L. Dooley, Rolf Kemler and Dietmar Vestweber for providing antibodies and Manuel Rebelo for providing B6/*nude* mice. We would like to thank Fábio Paiva and Faiza Al-Dalali for sample collection used in sequencing analyses, Ana C. Araújo, Maurícia Vinhas, Cláudia Florindo and André Mozes for technical assistance, and José A. Belo and Gabriela A. Silva for continuous support.

